# The parafascicular thalamus steers attention to facilitate learning

**DOI:** 10.64898/2026.06.18.733204

**Authors:** Louisa C. Kuper, Liliana M. Rohlf, Amy R. Wolff, Benjamin T. Saunders

**Affiliations:** Department of Neuroscience, University of Minnesota; Medical Discovery Team on Addiction, University of Minnesota; Graduate Program in Neuroscience, University of Minnesota

## Abstract

Cues that predict reward can acquire strong power to bias orientation and approach, directing seeking behaviors into close proximity with reward. The parafascicular thalamus (PF), part of the intralaminar complex, is well-placed as a neural hub for integrating subcortical sensory and attentional signals to generate actions that support cue-guided behavior. Neurons in the PF respond to sensory cues and encode features of head position, but little is known about how the PF is engaged in vivo during learning. Here, we recorded calcium activity in PF neurons using fiber photometry throughout a pavlovian conditioning task. We found that PF neurons developed sustained cue-evoked responses that scaled with associative learning and diminished with extinction. PF neurons preferentially signaled cue-directed orientations and body movements made during the cue, and their activity paused during reward consumption. Using optogenetics, we found that stimulation of PF neurons disrupted normal cue-directed orientation and impaired associative learning by promoting exaggerated ipsilateral turning behavior. Broadly, these results suggest an encoding profile by which the PF can support cue and reward approach behavior via dynamic regulation of head and body position. Together, our data demonstrate a thalamic region that is important for steering attention to facilitate cue-reward learning.

## INTRODUCTION

As animals navigate their environment, they must direct attention towards salient stimuli to learn about their identity, location, and relationship with rewards or threats. Cues that predict reward can acquire strong power to bias orientation and approach, directing seeking behaviors during times that reward is most likely available. For this process to be successful, sensory and motor signals must be integrated with brain centers for associative learning, motivation, and decision making (Grima et al., 2025; Klaus et al., 2019; Makino et al., 2016; Saunders & Robinson, 2013).

The parafascicular (PF) thalamus is an intralaminar nucleus that is well-placed to be a key modulator of this cued-guided behavior. It receives sensorimotor information from the superior colliculus (Krout et al., 2001; Lee et al., 2020; Redgrave et al., 2010; Schulz et al., 2009), frontal eye fields (Benarroch, 2023), and cerebellum (Deniau et al., 1992; Hintzen et al., 2018), as well as brainstem inputs related to arousal (Barroso-Chinea et al., 2011; Huerta-Ocampo et al., 2020; Yan et al., 2008). It sends dense excitatory output to the striatum (Berendse & Groenewegen, 1990; Elena Erro et al., 2002; Foster et al., 2021; Gonzalo-Martín et al., 2024; Mandelbaum et al., 2019), a region classically associated with cue-reward learning (Collins & Saunders, 2020; Cox & Witten, 2019). The PF also has reciprocal cortical connectivity as well as dense inhibitory input from the substantia nigra, and thus is set up to be an important interface between systems integrating attention with learning and action (Alloway et al., 2014; Gonzalo-Martín et al., 2024; Grillner, 2025; Lee et al., 2020; Mandelbaum et al., 2019; Redgrave et al., 2010; Schulz et al., 2009; Stayte et al., 2021). The PF is known to respond to salient sensory events (Matsumoto et al., 2001; Sicre et al., 2024), and is involved in action initiation (Díaz-Hernández et al., 2018; Wolff et al., 2022) and flexible decision making (Bradfield & Balleine, 2017; Brown et al., 2010; Kato et al., 2011). PF neurons are known to encode detailed features of head position, and stimulation of their activity generates head movements (Fallon et al., 2023; Watson et al., 2021). Despite these findings, we know little about how PF neurons are engaged during cue-guided learning and reward seeking.

To investigate this, we used fiber photometry to record population-level calcium signals in PF neurons while freely moving rats engaged in an associative learning task where a discrete cue predicted sucrose delivery (Flagel et al., 2011; Saunders & Robinson, 2012). We found that PF neurons were strongly activated by this reward predictive cue, in a manner that scaled with acquisition of conditioned approach behavior and decayed in extinction. As learning progressed, PF neurons showed sustained activity during cue periods, and preferentially encoded cue-elicited movement and orientation behaviors directed at the cue/reward location.

When rats paused to engage in reward consumption, we saw a concurrent dip in PF neuron activity. Finally, we found that optogenetic activation of neurons in the PF disrupted normal cue-directed orientation and impaired associative learning by promoting exaggerated ipsilateral turning behavior. Broadly, these results suggest an encoding profile by which the PF can support cue and reward approach behavior via dynamic regulation of head and body position within a given context. These results highlight a thalamic node that is important for steering actions to ensure environmental stimuli receive the attention that is required for learning.

## RESULTS

### PF neurons are activated by reward predictive cues and track associative learning

To investigate the role of the PF thalamus in cue-directed learning, we expressed the calcium indicator GCaMP8f in PF neurons (Fig 1A-C) and used fiber photometry to record population activity during a pavlovian conditioning task (Fig 1D). During training, the cue, a lever, was extended for 8 seconds, then retracted, which coincided with a liquid sucrose reward delivered to an adjacent port, with trials separated by a variable interval (Fig 1D). We measured behavior in two primary ways. A DeepLabCut (DLC) pipeline was used to quantify rat’s position and movement throughout the chamber, and port entries were logged by the Med Associates equipment. Rats learned the lever cue-reward association over 12 days of training, engaging in physical approach to the cue/port area and spending significantly more time in the reward port during cue presentations (Fig 1E-F, main effect of session, F(1.761,35.23) = 26.52, p>0.0001). Early in training, PF neurons showed a robust increase in activity at the onset of the cue (Fig 1G,H). As training progressed, this initial signal peak was maintained, but a sustained increase in activity developed during the entire cue period, as measured by area under the curve (AUC, Fig 1H-I, 2 way RM ANOVA, main effect of session F(3,33) = 6.455, p=0.0015). We saw no change in activity in PF neurons when signals were aligned to control windows during the intertrial interval (Supp. Fig 1D). Further, control recordings made from rats with a YFP-only virus expressed in the PF showed no cue-related signal changes (Supp. Fig 1E).

**Fig 1.**
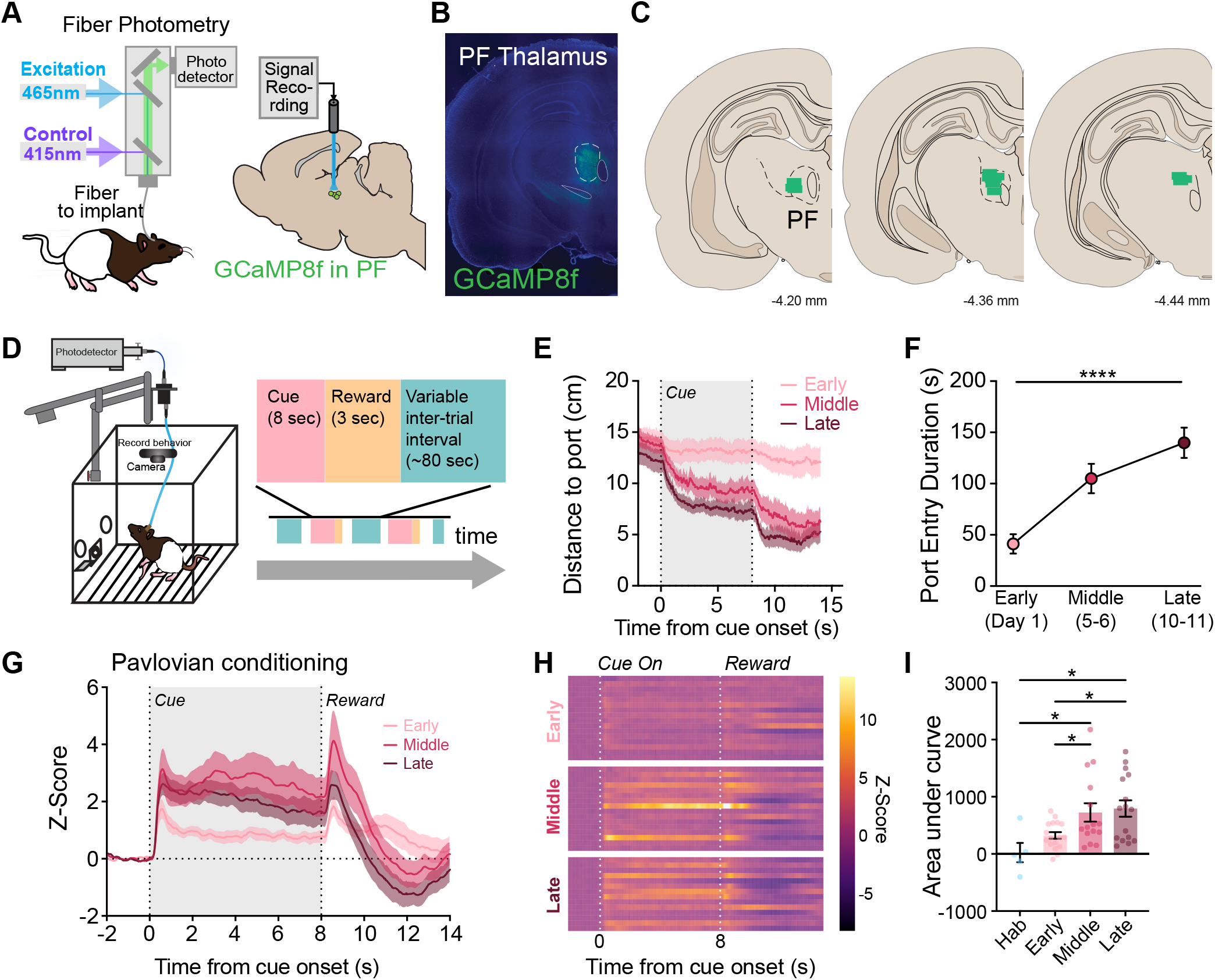
PF neurons track associative learning. The calcium indicator GCaMP8f was expressed in the parafasicicular (PF) thalamus for in vivo fiber photometry recordings of neural activity in rats (n=19). B) Representative image of GCaMP expression in the PF. C) Recording fiber placements in the PF. D) Behavioral paradigm for lever-reward conditioning. The conditioned stimulus, a lever, was extended for 8 seconds, then retracted, which coincided with a liquid sucrose reward delivered to the adjacent port. Cue-reward pairings were separated by a variable intertrial interval averaging 100 seconds. E) The lever cue evoked approach behavior, reflected by an increased proximity to the reward port at cue onset across learning. F) Rats spent significantly more time in the reward port during the cue across early (Day 1), middle (Days 5/6), and late (Days 10/11) phases of conditioning (p>0.0001). G) Z-scored traces of calcium recordings of PF neurons aligned to cue onset across learning phases. PF neurons rapidly developed a response to cues, which increased as learning progressed. H) Heatmap of cue responses across individual days for all animals during early, middle, and late phases of training. I) Neural activity during the cue period significantly increased across training, as measured by area under the curve (AUC, p=0.0015). Data shown reflect mean +/-SEM. *p<.05, ****p<.0001.

### Sustained cue-evoked PF neuron activity diminishes during extinction

After conditioning, rats underwent extinction in sessions where the lever cue was presented without the sucrose reward (Fig. 2A). Rats quickly updated their behavior, no longer approaching the reward port area and significantly decreasing their reward port entry duration during cue presentations (Fig. 2B,C, Mixed-effects analysis, F(1.020,21.43) = 69.23, p<0.0001). Relative to the large sustained signals seen late in reward conditioning (Fig. 2D), cue-related PF neuron activity diminished during extinction (Fig. 2E,F) with significantly lower area under curve during the cue period (Fig 2G, 2-way ANOVA main effect of session, F(3,22) = 5.880, p=0.0042).

**Fig 2.**
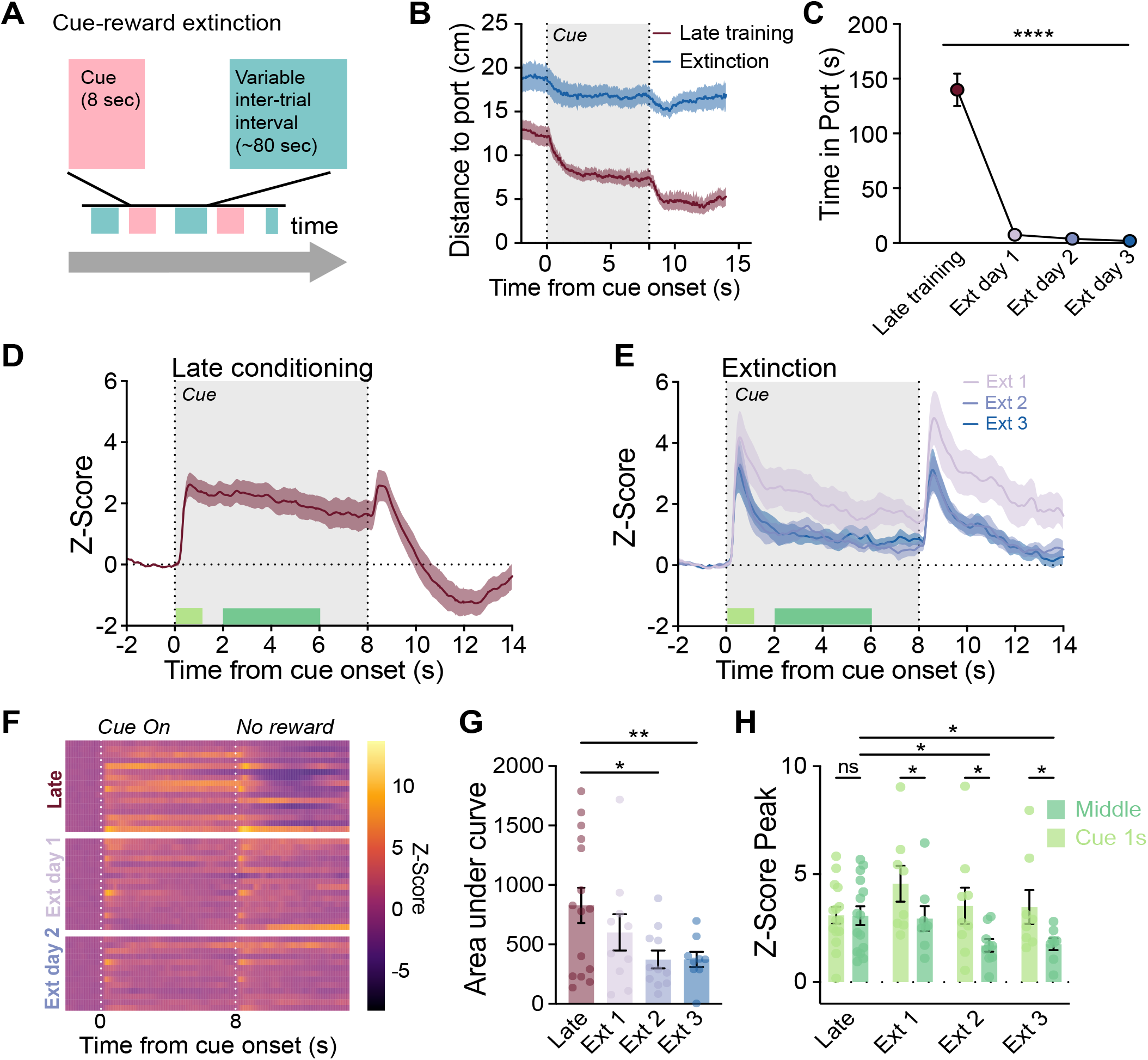
Sustained reward cue-evoked PF neuron activity diminishes during extinction. A) Schematic of extinction behavior paradigm. Rats (n=10) were exposed to the same cues and inter-trial intervals, but with reward omitted, for 3 sessions. B) Cue-evoked approach behavior decreased in extinction. C) Extinction resulted in a rapid decline in reward port duration during the cue period (p>0.0001). D) During the late phase of reward conditioning, PF neurons showed sustained activity during the cue period. E) Across extinction, PF activity at cue onset and offset remained high, but the sustained activity throughout the cue diminished. F) Heatmap of cue responses during late conditioning versus extinction days. G) Signal AUC during the cue period significantly decreased across extinction (p=0.0039). H) Signal peaks during cue onset (0-1.5 sec) vs the middle of the cue period (2-6 sec). During extinction, peak activity in the middle period was significantly lower than at cue onset (Mixed effects analysis, effect of cue window, F(1,14) = 7.95, p=0.01) during all days of extinction. The middle of the cue period significantly decreased from late training during extinction (Mixed effects analysis, effect of day F(3,42) = 3.30, p=0.0293), with no significant changes in the first second after the cue period. Data shown reflect mean +/-SEM. *p<.05, **p<.01, ****p<.0001.

Notably, a large increase in PF activity remained at cue onset and offset during extinction (Fig. 2E,F). To measure differences in signal throughout the cue period, we compared the peak Z-Score during the first 1.5 sec of the cue to the peak across the middle (2-6 sec) period of the cue (Fig. 2D,E,H). In late training, there was no significant difference between these periods, indicating a sustained response (Fig. 2H). In contrast, during extinction, there was a significant drop in peak activity between cue onset and middle periods (Fig 2H, Mixed effects analysis, main effect of window, F(1,14) = 7.95, p=0.0136). Together with Figure 1, these data demonstrate that PF neurons track the emergence and decay of pavlovian cue-reward associations.

### PF neurons preferentially signal cue-evoked movements

As rats learn cue-reward associations, the cue comes to drive expression of more vigorous movement, in this case directed at the cue and reward location. To isolate movement-related components of PF neuron activity in our task, we compared cue period signals to control periods. The control windows were 8-second periods during the intertrial interval, ∼15 seconds before each cue presentation (Fig. 3A). To investigate movement features such as locomotion state and speed, we made use of our DLC pipeline (Fig. 3B-C). Looking at the entire 8-sec period for cue and control windows, there were no overall differences in movement metrics, including locomotion bouts and speed (Fig. 3D-F, Average speed: Mixed-effects analysis F(1,16) = 0.00062, p=0.9804), Percent locomotion bouts: Mixed-effects analysis, F(1,16) = 0.2707, p=0.6100), Duration of locomotion bouts: Mixed-effects analysis, F(1,16) = 0.06710, p=0.7989).

**Fig 3.**
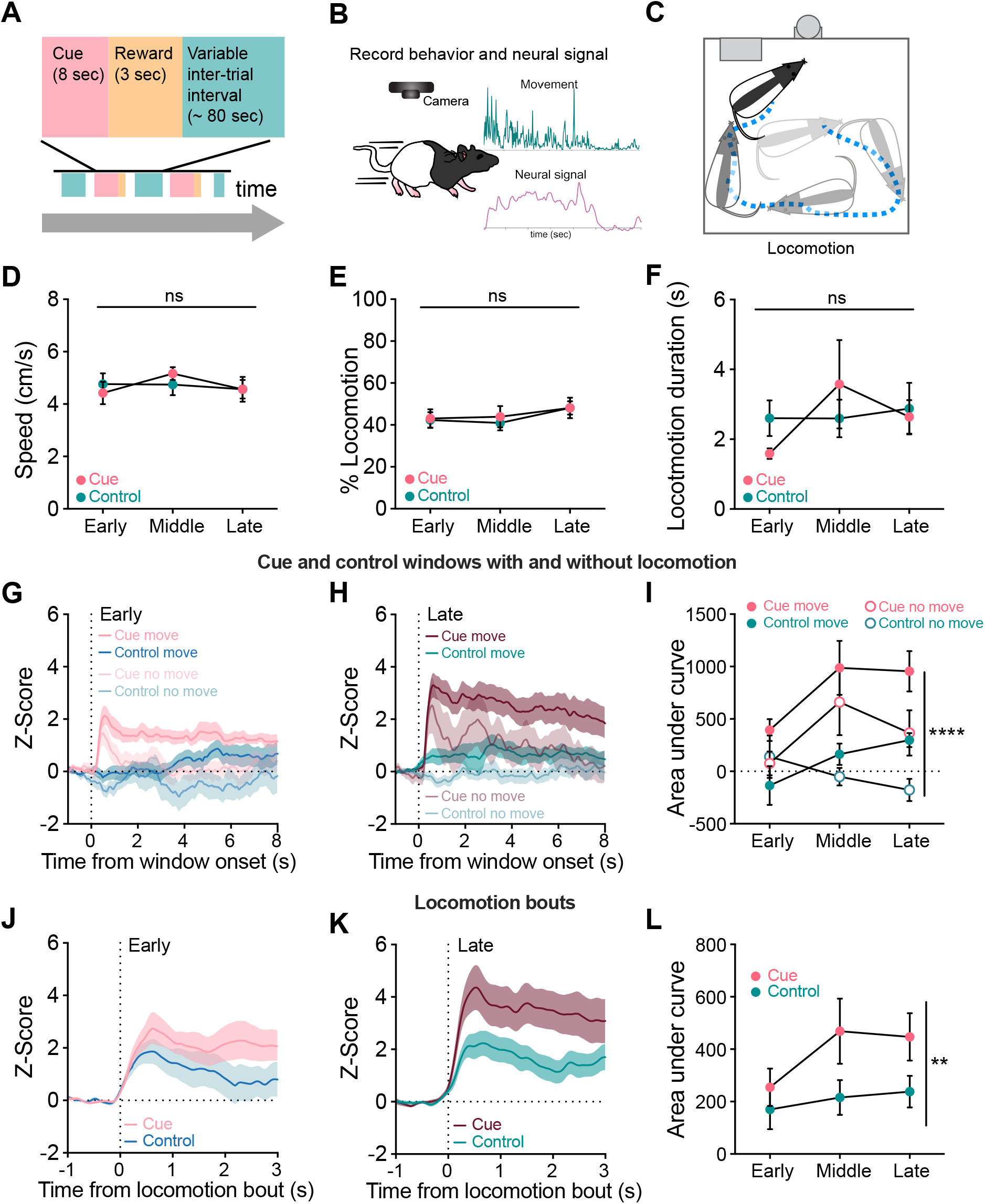
PF neurons preferentially signal cue-evoked movements. A) Schematic of pavlovian behavioral paradigm. B) DeepLabCut pipeline. Recordings were aligned to body part movements to assess speed and locomotion. C) Locomotion was defined as a bout of a minimum 0.5 seconds, with a threshold of 0.3 seconds of no movement to be considered the start/end of a bout. Cue periods were compared to control periods taken from a random 8 second window within the inter-trial interval. D) There was no difference in the average speed during the cue and control periods. E) There was no difference in percent locomotion bouts detected between cue and control periods. F) There was no difference in the duration of locomotion between cue and control periods. G) PF activity aligned to the beginning of cue or control windows when locomotion occurred during early training. H) PF activity aligned to the beginning of cue or control windows when locomotion occurred during late training. I) Signal AUC for cue and control windows with versus without locomotion. PF activity was significantly different between conditions (p<0.001) and highest when movements occurred during the cue. J) PF activity aligned to the onset of locomotion bouts in cue and control periods during early training. K) PF activity aligned to the onset of locomotion bouts in cue and control windows during late training. L) Signal AUC for cue and control locomotion bouts across training. PF activity was significantly higher during cued locomotion bouts (p=0.0287). Data shown reflect mean +/-SEM. **p<.01, ****p<.0001.

We next sorted cue and control period PF neuron signals based on whether or not a locomotion bout occurred in each window category. In early training, there was a moderate increase in cue period PF activity on trials where there was locomotion, and minimal increase from baseline in control periods with locomotion (Fig. 3G). By late training, despite no difference in average speed, cue periods where locomotion occurred had greater PF activity compared to control periods with locomotion (Fig. 3H,I; Mixed-effects analysis main effect of window, F(3, 48) = 10.73, p<0.001). PF activity was modulated by locomotion state, as within both the cue and control periods, there was greater PF activity on trials where locomotion occurred (Fig. 3I). Critically, however, even on trials where no locomotion occurred, we saw phasic increase in PF neuron activity at cue onset (Fig. 3. G,H).

We next z-scored PF signals to locomotion onset within each window period. Both cue and control window bouts of locomotion resulted in an increase in PF activ-ity in early training (Fig. 3J). By late training, however, cue-related locomotion bouts were characterized by significantly larger signals than control window bouts (Fig. 3K,L, Mixed-effects analysis, F(1,16) = 9.90, p=0.0062). Overall, these data demonstrate that PF neurons are modulated by movement state in a way that intersects with associative learning, to preferentially signal cue-evoked locomotion.

### PF neurons preferentially encode cue-evoked orientations

Given that we saw greater PF activity during cue periods, which grew concomitant with increased conditioned behavior, we speculated that PF signals may be related to the development of vigorous cue/reward-directed movement. Using our DLC pipeline, we defined an orientation as a period when a vector from the rat’s implant to mid back was within 30 degrees of the cue/reward zone during either the cue period or the control windows (Fig. 4A). To quantify the emergence of cue/reward directed orientation, we calculated the percent of time spent orienting to different positions in the chamber visualized as polar plots (Fig. 4B,C). Across training during the cue period, rats developed an orientation bias, spending more time with their head pointed to the cue/reward portion of the chamber (Early cue vs. late cue Kolmogorov-Smirnov test, D=0.4722, p=0.0007). This orientation bias was specific to only when the cue was present, as we did not see a differentiation of orientation to the cue/reward zone during the control windows across training (Early control vs. late control K-S test, D= 0.1944, p=0.5041). We next Z-scored PF signals to the onset of orientations. In early training, there was little difference in PF activity during cue-evoked orientations and orientations made during the control window (Fig. 4D). By late training, however, cue-evoked orientations resulted in significantly greater PF activity compared to control window orientations (Fig. 4E,F; AUC, Mixed-effects analysis main effect of window F(1,16) = 21.10, p=0.0003). Taken together with our data in Figure 3, these analyses show that PF population activity is correlated with head position in a way that preferentially encodes orientations to important stimuli in the environment.

**Fig 4.**
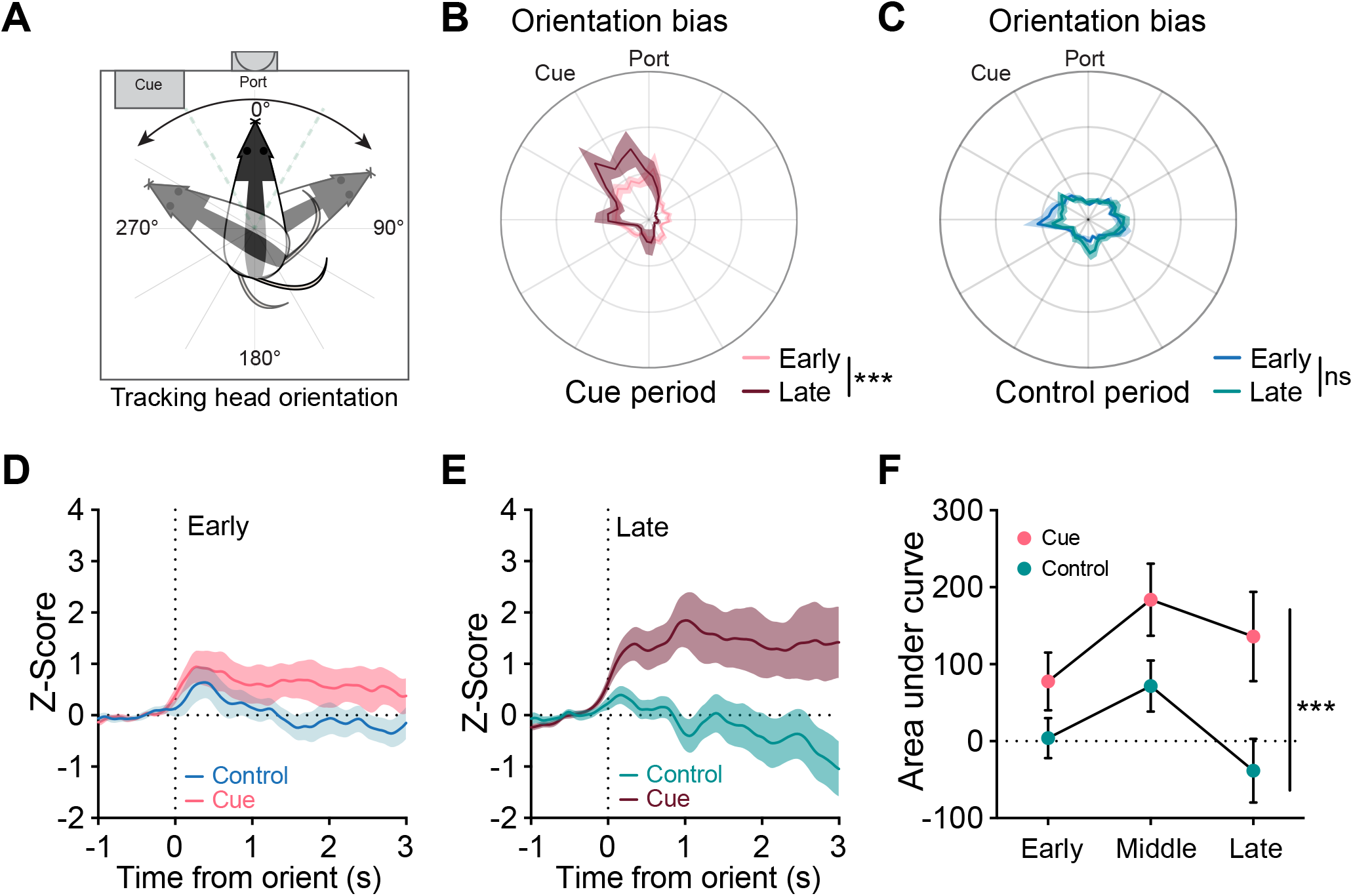
PF neurons preferentially encode cue-evoked orientations. A) Orientations were defined as the period when a vector from the rat’s implant to nose tip was within 30 degrees or less of the reward magazine for a minimum of 0.4 seconds during either the cue period or the control period. B) Polar plots of orientation during cue periods early versus late in training show the emergence of bias for head direction aimed at the cue/port area (Early cue vs. late cue K-S test, p=0.007, Early control vs. late control K-S test, p=0.541). C) Polar plot of orientation during control windows early versus late in training shows no such bias. D) PF activity aligned to onset of orientations made during the cue or control window during early E) or late training. F) Area under curve of cue vs control window orientations. Activity during cue-related orientations was significantly higher than for control-window orientations (p=0.0003). Data shown reflect mean +/-SEM. ***p<.001.

### PF neurons pause during reward consumption

Little is known about how reward consumption engages PF neuron activity. In our task, reward delivery was signaled by the retraction of the lever cue. Examining PF activity averages, we saw an increase in PF signal co-incident with lever retraction/reward delivery onset, followed by a dip in signal below baseline (Fig 1). Rewards were not always consumed exactly at cue offset, and trained rats often entered the port during the cue and stayed there until reward was given. To separate the contribution of these nuances in behavior to PF neural signal, port entries were sorted into three types: port entries that occurred during the cue (Cue PE), port entries in which reward was consumed (Consume PE, defined by spending at least 0.5 sec within the port after reward was delivered to ensure rats were drinking sucrose), and port entries during the ITI (Fig. 5A).

**Fig 5.**
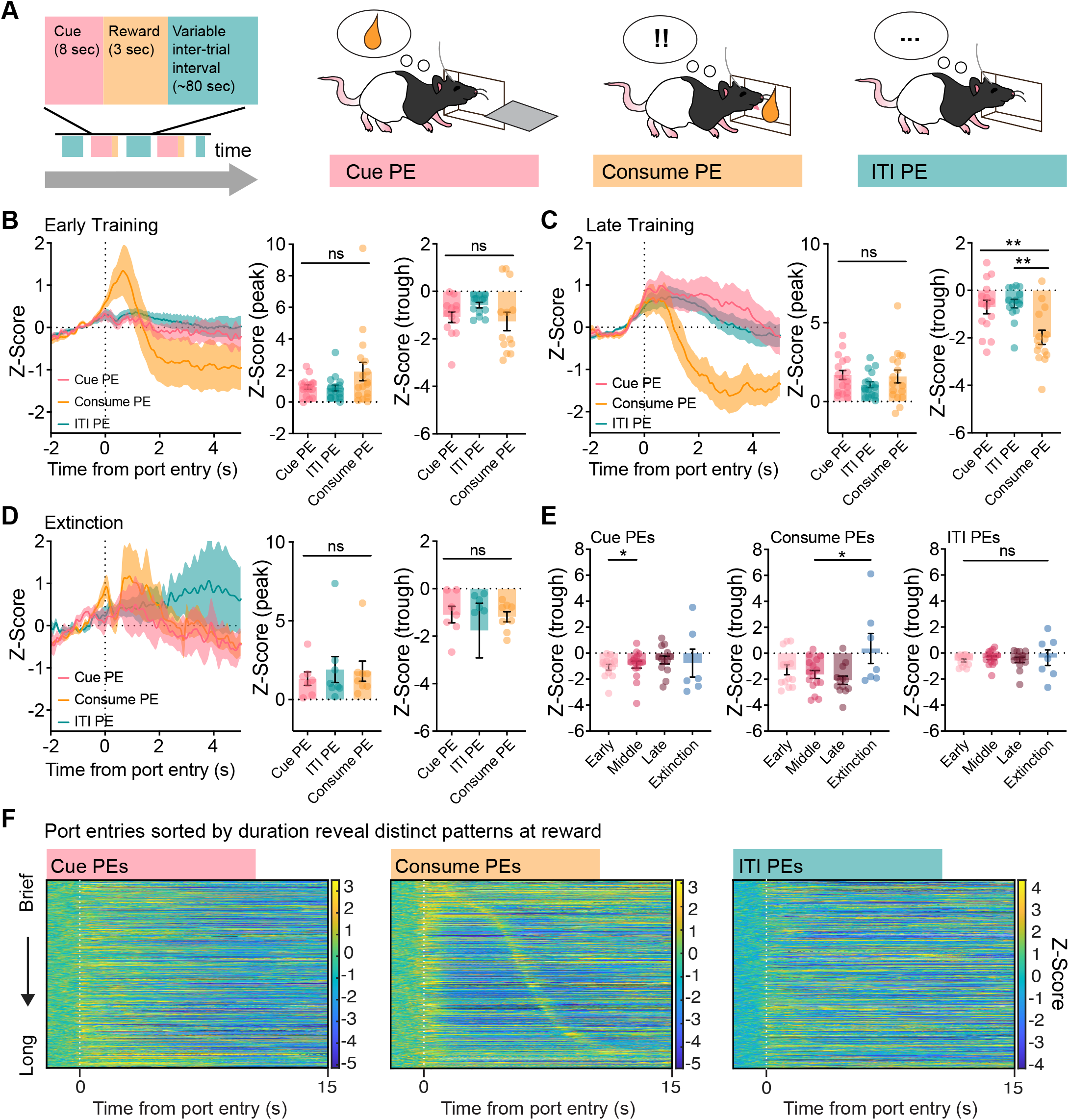
PF neurons pause during reward consumption. A) Port entries (PE) were separated according to three time windows: the cue period, the reward consumption period, and the inter-trial interval (ITI). B) PF neuron activity aligned to each port entry type early in training. There was no difference in signal peak or trough across PE types. C) PF neuron activity aligned to each port entry type late in training. Consumption port entries had significantly lower trough than cue or ITI port entries (p=0.0005). D) PF neuron activity aligned to port entry types during extinction. There were no differences in signal peak or trough. E) Signal trough for each port entry type across experimental phases. The negative signal associated with consumption port entries increased across conditioning and then went away during extinction (p=0.0026). F) Heatmaps of all trials of each port entry type sorted by their duration (brief to long top to bottom) in the late training phase. During consumption PEs, PF neuron activity dipped for the duration of the consumption event. Data shown reflect mean +/-SEM. **p<.01, **p<.01.

To isolate how PF neurons responded to reward consumption, we aligned PF calcium signals to port entries of each type. As training progressed, consumption port entries were characterized by a distinct biphasic activity pattern, with a positive peak followed by a dip in signal below baseline. Cue and inter-trial interval port entries, by comparison, were characterized by a simple moderate increase in PF activity (Fig. 5B,C).

Given the biphasic nature of the signals, we plotted the Z-scored signal peak (maximum) and trough (minimum) during port entries, separated by type. Peak and trough signal values were similar across port entries types early in training (Fig. 5B, trough One-way ANOVA F(2,48)= 2.512, p=0.0917, peak one-way ANOVA F(2,48) = 2.640, p=0.0817). By late training peak signal did not vary by port entry type (Fig. 5C, One way ANOVA F(2,48) = 1.144, p=0.3271), but the signal trough was significantly lower for consumption port entries relative to port entries made during the cue or the ITI (Fig. 5C, F(2,42) = 9.164, p=0.0005, Consume PE vs. Cue PE p = 0.0031, Consume PE vs ITI PE p=0.001).

We next investigated how port entry responses evolved during extinction, when port entry behavior was rapidly diminishing (Fig. 3B). We analyzed extinction port entries types in the same manner to those during conditioning with reward, but with no consumption threshold. As extinction progressed, there was no change in the peak of PF signals (Fig 5D, One-way ANOVA F (2,20) = 0.2067, p=0.8150), but the consistent dip in PF activity during “consumption” port entries was eliminated (Fig. 5D). Comparing how all port entries evolved across time, only “consumption” port entries were affected by extinction, with significantly smaller trough values (Fig. 5E, mixed-effects analysis effect of day, F (2.236,21.61) = 7.528, p=0.0026).

To clarify whether drops in PF activity matched with rat’s port entry engagement, we sorted port entry signals during reward conditioning by duration (Fig. 5F). Only consumption port entries showed a clear relationship with PF signals based on their duration. PF neuron activity had an initial increase at port entry onset, followed by a dip in signal throughout consumption (duration of port entry), with an increase at consumption offset (Fig. 5F, middle).

Overall, the patterns of activity from our fiber photometry recording studies suggest that PF neurons are engaged as rats orient attention to a learned cue/reward location, and their activity pauses when rats are inactive during reward consumption.

### Disrupting endogenous PF thalamus activity impairs cue-directed attention and learning

So far we have demonstrated that PF neurons are engaged at the population level during associative learning, encoding a conditioned stimulus and preferentially signalling cue/reward-aimed movements. Broadly, this suggests that phasic PF signals are important for directing attentional shifts and exploration of important sensory elements in the environment. Based on this, we hypothesized that disruption of endogenous PF activity may impact the normal progression of associative learning.

To investigate this, we expressed the excitatory opsin ChR2 in PF neurons and implanted fibers for optogenetic modulation (Fig. 6A,B). Rats (n=12) were put through the same lever-sucrose conditioning paradigm as above. Laser pulses, to excite PF neurons, were administered on a subset of training days (Days 1, 5, 9, and 12). On stimulation sessions, laser delivery (5 sec, 20 Hz) coincided with cue onset (paired group, n=8) or during a period of the ITI (unpaired group, n=4), on 50% of trials (Fig. 6C,D).

**Fig 6.**
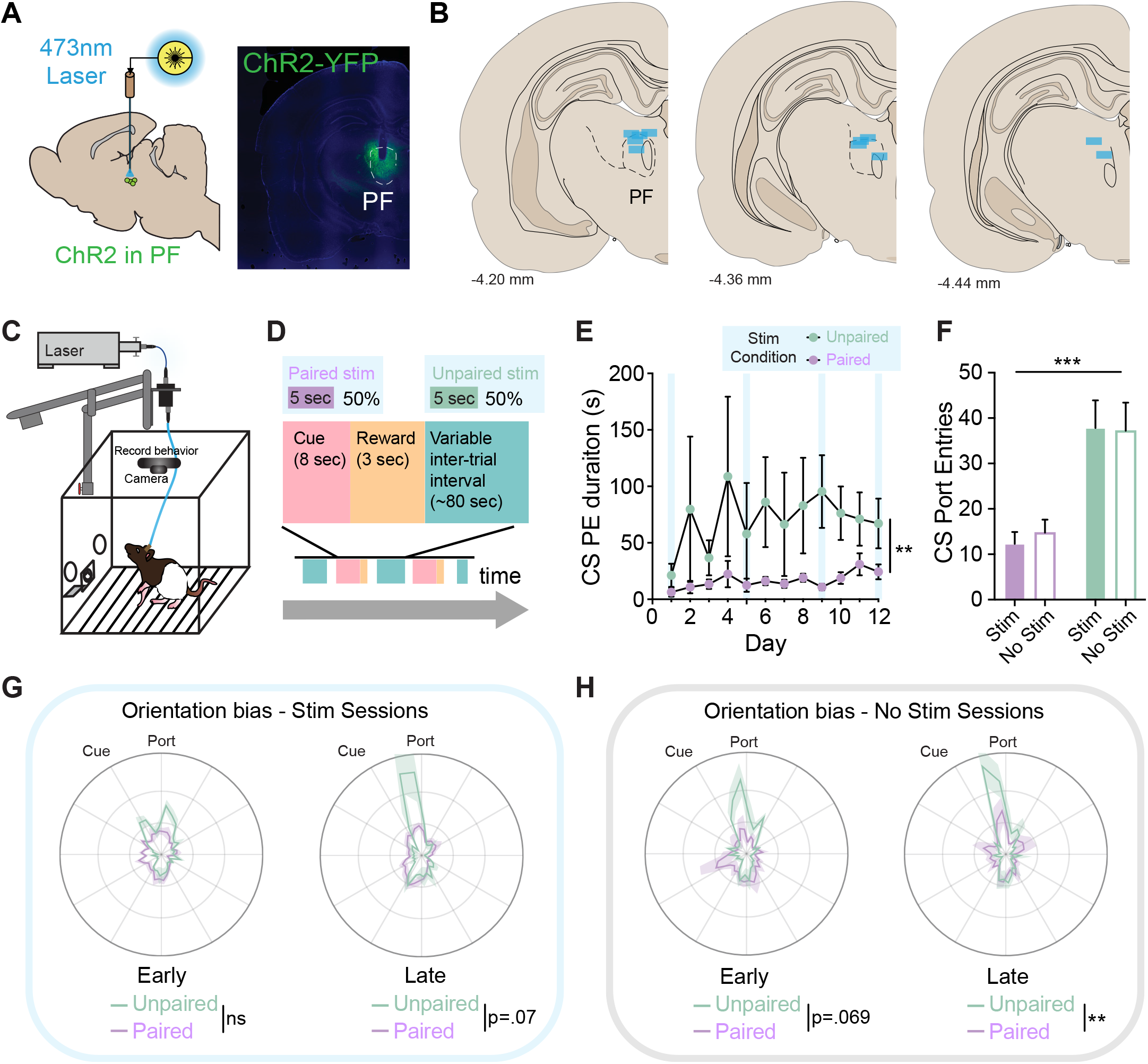
Disrupting endogenous PF neuron activity impairs cue-directed attention and learning. A) ChR2-YFP was expressed and optic fibers were implanted unilaterally into the PF thalamus. B) Summary of optic fiber placements in the PF. C) Rats were trained on a pavlovian conditioning task where they learned to associate a lever with sucrose. D) Laser stimulation (20 Hz, 5 s) was delivered on a subset of training days (1, 5, 9, 12) either during the cue period (paired group, n=8) or inter-trial-interval (unpaired group, n=4). E) Across training, paired rats spent significantly less time in the port during the cue period compared to unpaired rats (Mixed-effects analysis, effect of stim condition, F(1, 10) = 11.12, p=0.0076). F) Port entries made during the cue were significantly lower on both stim and no stim days for paired rats compared to unpaired rats (F(1,10) = 18.93, p=0.0014). G) Polar plots of orientation bias (mean proportion of time at angle) during cue periods for Paired vs. Unpaired rats on early and late stim sessions (Early, K-S test p=0.1243, Late K-S test p=0.0694). H) Polar plots of orientation bias during cue period during no stim sessions (Early K-S test p=0.0694, Late K-S test p=0.0039). Data shown reflect mean +/-SEM. **p<.01.

As in our earlier experiments, we examined the development of conditioned port entry behavior. We found that excitation of PF neurons profoundly impaired performance on the task. Across training, while time in the port increased overall (Fig. 6E; Mixed-effects analysis main effect of training F(11,107) = 7.672, p<0.0001), paired stimulation rats had an impaired learning trajectory (Mixed effects analysis session x stim condition interaction, F(11,107) = 2.138, p=0.0234), spending significantly less time in the port during the cue period compared to unpaired rats (Fig. 6E, effect of stim condition, F(1,10) = 11.12, p=0.0076). This was true on the stimulation days, but the effect persisted in sessions where no laser was delivered for either group (Fig. 6F, F(1,10) = 18.93, p=0.0014), indicating that paired stimulation disrupted the formation of the cue-reward association, altering the trajectory of conditioned responding.

We next examined how rats’ orientation to the reward port was impacted by optogenetic activation of the PF. As above (Fig. 4), using body angle vectors from our DLC model, we calculated the proportion of time rats were oriented to the reward port during cue periods, and mapped those values as a polar plot of angles (Fig. 6G,H). This analysis revealed that optogenetic activation of PF neurons paired with the cue prevented the development of an orientation bias to the cue/reward location of the chamber, as compared to the unpaired stimulation group (Fig. 6G; Early K-S test D=0.2778, p=0.1243; Late K-S test D=0.3056, p=0.0694). This effect was also evident on the sessions where no laser stimulation was delivered (Fig. 6H; Early K-S test D=0.3056, p=0.0694; Late K-S test D=0.4167, p = 0.0039), again demonstrating that PF activation impaired learning.

### PF stimulation potentiates ipsiversive turning

Given the dysregulation of cue-directed attention and conditioned behavior we saw with optogenetic manipulation of PF neurons (Fig. 6), we examined the behavioral impact of stimulation in the paired rats more closely, making use of our DLC pipeline to quantify detailed features of movement (Fig. 7A-C). We first calculated movement speed during the cue period on early (day 1) and late (day 12) days of conditioning, for trials where laser was delivered (stim) and those without laser (no stim). For no stim trials, speed rapidly increased at cue onset and then quickly returned to baseline (Fig. 7D,E). For stim trials, this movement was more sustained, such that speed values were larger and elevated speed persisted for the entire laser delivery window, decreasing at laser offset (Fig. 7E-G; 2 way RM ANOVA, main effect of laser, F(1,7) = 8.158, p=0.0245). Average speed on stimulation trials was significantly higher than on trials without stimulation (Fig. 7G, 2 way RM ANOVA, main effect of laser, F(1,7) = 8.158, p=0.0245). The impact of stimulation on movement potentiated over time, such that speed on stim trials late in conditioning was higher compared to the early conditioning time point (Fig. 7G, 2 way RM ANOVA, main effect of session, F(1, 7) = 21.80, p=0.0023).

**Fig 7.**
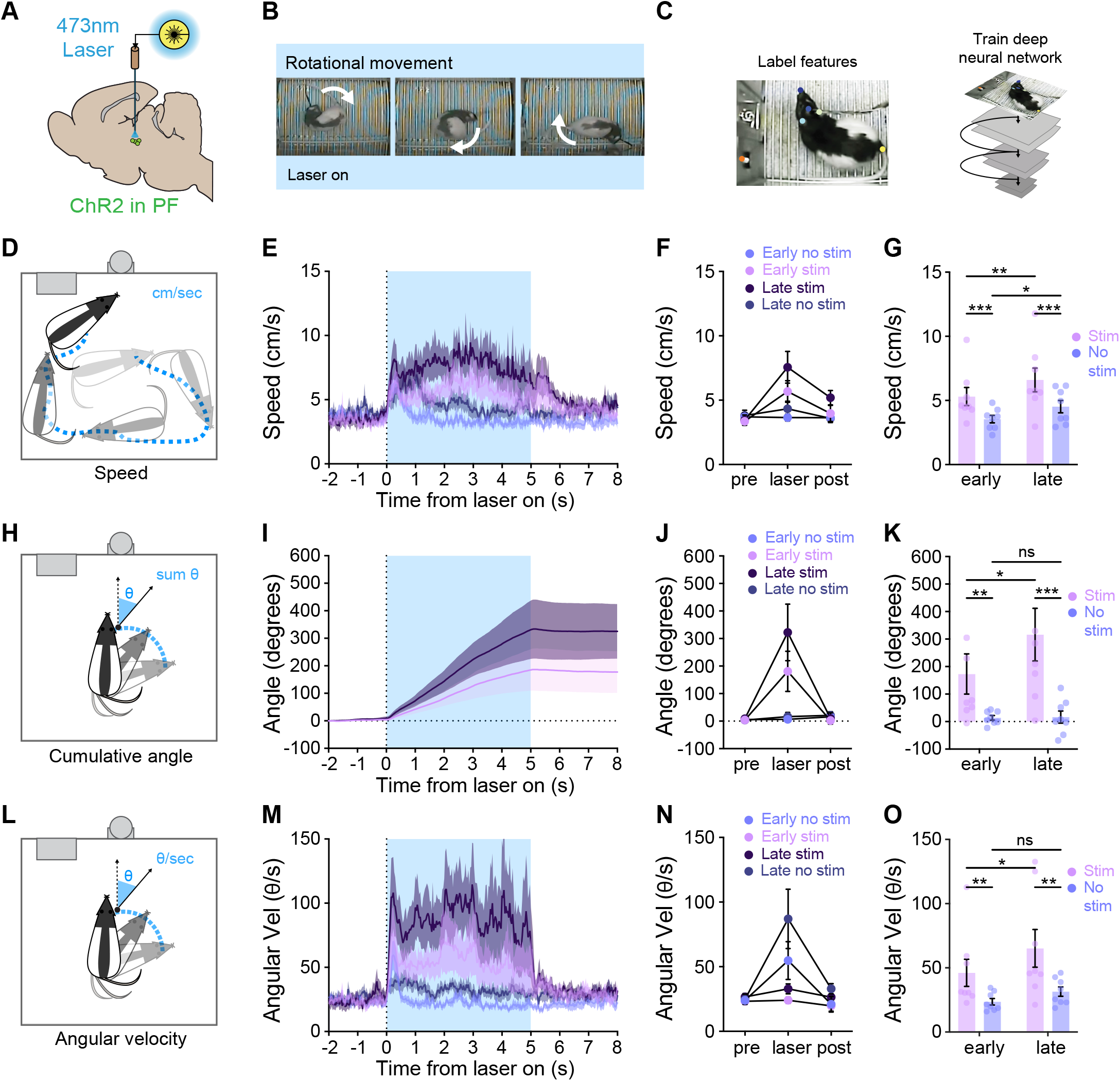
PF stimulation potentiates ipsiversive turning. A) ChR2 was injected into the PF unilaterally with ChR2 (n=8). B) Representative images of rotational movement while the laser was on. Rats curled and turned their body in circles, led by the head. C) Schematic of behavioral analysis via DeepLabCut. D) Representative image of speed calculations. Speed was calculated as rats’ distance traveled over time in cm/sec. E) Average speed traces over the laser period (shaded in blue). F) Speed increased over the laser period. G) Average speed during cue period was higher on laser paired trials (p=0.0245). Average speed increased in training for both stimulation and no stimulation trials (p=0.0023), but increased to a greater degree in laser paired trials (p=0.0140). H) Representative image of cumulative angle calculations. Cumulative angle was measured as the sum of the number of degrees a vector from their shoulder to implant shifted across the cue period. I) Cumulative angle traces during the cue period. During stimulation trials, cumulative angle increased throughout the laser period. Cumulative angle had very low increases for trials without stimulation and control (YFP) animals during stimulation. J) Cumulative angle increased exclusively during the laser period. K) In both early and late training, cumulative angle was significantly higher on stimulation trials versus trials without stimulation (p=0.0258). Average cumulative angle increased across learning for stimulation trials only (p=0.0161). L) Representative image of angular velocity calculations. Angular velocity was calculated as changes in absolute body angle in degrees per second. M) Traces of average absolute angular velocity. N) Angular velocity during the cue period was significantly higher on stimulation trials, and had little change on trials without stimulation. O) In both early and late training, angular velocity was significantly higher on stimulation trials versus trials without stimulation (p=0.0365). Average angular velocity increased across learning for stimulation trials only (p = 0.0188). Data shown reflect mean +/-SEM. *p<.05, **p<.01, ***p<.001.

From visual inspection of behavior videos it was evident that movement evoked by optogenetic activation of the PF was not linear, forward-directed body repositioning. Instead, the predominant effect of PF excitation was turning behavior (Fig. 7A-C). We assessed this rotational movement by quantifying head angle deflection, summing change in position of a vector from their shoulder to implant across the cue period (Fig. 7H). Angles were normalized based on the injection site of the rat, as rats uniformly turned ipsilaterally to their injection hemisphere. Changes in cumulative angle were tightly locked to laser stimulation, increasing in a linear fashion throughout the laser period, and ceased to change once the laser was turned off (Fig. 7I, J). Cumulative angle was significantly higher on stimulation trials compared to trials without stimulation (Fig. 7I-K, 2 way RM ANOVA, main effect of treatment, F(1,7) = 7.946, p=0.0258), where no sustained head deflections were seen. Cumulative angle significantly increased from early to late sessions for stimulation trials only (Fig. 7I-K, paired t-test, stim early vs late t(7) = 3.150, p=0.0161).

We hypothesized that the overall increase in speed we observed (Fig. 7E-G) reflected a change in the speed of head turning, given the rotational nature of the movement. We quantified this as angular velocity, based on angle of turning in degrees per second (Fig. 7L). Cue onset resulted in an increase in angular velocity, which was larger and more sustained on laser stimulation trials (Fig. 7M, N). Trials with stimulation had a significantly greater average angular velocity compared to no stimulation trials (Fig. 7N,O; 2 way RM ANOVA, main effect of treatment, F(1,7) = 6.651, p=0.0365). Angular velocity also increased from early to late sessions for stimulation trials only (Fig, 1O, paired t-test early vs late t(7) = 3.041, p=0.0188).

We quantified movement in the unpaired conditioning rats, which received laser stimulation during the ITI of the cue-reward conditioning paradigm (Suppl Fig 2). These rats also showed ipsilateral head/body turning that was time-locked to laser stimulation, as measured by angular velocity (Suppl Fig. 2D). We found no consistent significant differences in turning measures for paired vs unpaired rats (Suppl Fig. 2E).

These results indicate that an important function of the parafascicular thalamus is to steer head and body orientation, consistent with some past studies (Fallon et al., 2023; Watson et al., 2021), but this function is potentiated when PF neurons are repeatedly active. Taken together with our data in Figure 6, our optogenetic studies show that exaggeration of normal PF activity impairs associative learning by interrupting directed attention required to link a cue with reward within a specific sensory context.

### PF neuron stimulation is not reinforcing

Given that the PF sends outputs to reward centers such as the nucleus accumbens (Berendse & Groenewegen, 1990; Y. Zhang et al., 2022), in a final study, we investigated if activation of PF neurons promotes reinforcement. Rats were given the opportunity to self-stimulate the PF in an intracranial self-stimulation (ICSS) procedure. Over 3 1 hr sessions, rats could choose to poke in an active port (designated by three cue lights) that would result in a 1 second 20 Hz stim of the PF, or an inactive port that had no consequence (Suppl. Fig. 3). Nose poke behavior was low, and there was no difference between nose poke durations to active versus inactive ports (2 way ANOVA, no effect of condition F(1,10) = 1.215, p=0.261).

## DISCUSSION

Here, we investigated the role of the parafascicular (PF) thalamus in cue-guided behavior. Our main findings are as follows: 1) Population-level photometry recordings of calcium activity show that PF thalamus neurons are dynamically activated as rats reorient their head and body position in the environment. 2) Critically, these signals were not simply a readout of motor state, as PF activity scaled with associative learning and preferentially signaled behaviors that were directed at a predictive cue/reward location, subsequently decaying during extinction. 3) During reward consumption we saw a decrease in PF activity. 4) Finally, disruption of normal PF neuron signals, via optogenetic activation, impaired cue-reward learning by generating exaggerated head movements that overrode cue orientation. Collectively, our data reveal new insight into a thalamic region that is important for steering attention to facilitate exploration and targeted actions that support learning. These results offer insight into sensory, attentional, and movement-related brain systems that govern approach behavior and dynamic foraging decisions.

### The PF thalamus is a neural interface of attention and action

To navigate dynamic environments, many animals rely on external sensory information, which focuses attention within complex scenes, to facilitate approach toward rewards. Much investigation has implicated dopamine and basal ganglia systems in driving cue-evoked approach behavior (Cox & Witten, 2019; Iglesias et al., 2023; Nicola, 2007; Saunders et al., 2018), but it has been less clear how sensorimotor signals are integrated to organize this process. Our results suggest that the PF thalamus serves as an interface between attention and action, integrating egocentric information (e.g., head position control) with allocentric information (e.g., cue/reward locations) to guide survey of the environment that underlies learning. Our data add insight into the broader literature on brain systems encoding space and body position, including the head direction system of the anterior thalamus (Taube, 1995), a key relay within hippocampal memory systems to represent the body’s internal compass (Angelaki & Laurens, 2020; Cullen & Taube, 2017; Grieves & Jeffery, 2017). The PF thalamus is also sensitive to head position but, notably, unlike many other thalamic regions, it sends dense projections throughout the striatum (Berendse & Groenewegen, 1990; Elena Erro et al., 2002; Gonzalo-Martín et al., 2024; Mandelbaum et al., 2019), and is degraded in Parkinson’s (Henderson et al., 2000; Villalba et al., 2014), suggesting it has a major role within basal ganglia dopaminergic circuitry for sensorimotor learning.

In contrast to the anterior thalamus head direction system, our data, and previous studies (Fallon et al., 2023; Watson et al., 2021), suggests that the PF thalamus does not encode spatial position of the head per se, but has populations of neurons that are dynamically engaged to promote specific head and body movements. We observed that PF neural activity is related to movement but, critically, is modulated by cue-reward learning. PF activity matched anticipatory reward consumption actions, including port entries during the cue period, cue-specific locomotion bouts, and orientations to the cue/reward location. Our results further suggest that the tuning of PF neuron activity to head movements is not fixed, but changes as salient sensory cues and locations are learned. PF neurons were preferentially active when rats made orientations to the learned cue/reward location, and paused during reward consumption. We found that optogenetically stimulating the PF during reward-predictive cues prevented proper pavlovian learning, suggesting that an exact pattern of endogenous PF activity is necessary for steering attention required to form cue-reward associations. Our results are consistent with past studies indicating that the PF is involved in learning movement sequences (Díaz-Hernández et al., 2018; Wolff et al., 2022), including directional reaching (Sibener et al., 2025). Our results offer new insight, identifying the PF thalamus as an important brain region for steering directed attention toward salient stimuli in the environment, to facilitate reward learning. This adds context to other previous work reporting PF neurons responses to reward-related cues (Matsumoto et al., 2001; Sicre et al., 2024; Yamanaka et al., 2018).

### Input-output circuit motifs for diverse PF thalamus functions

Our results support the notion that the PF thalamus sits at the intersection of circuits that support exploration and learning. The PF receives sensorimotor information from the superior colliculus and cerebellum, arousal signals from the brainstem, action signals from the substantia nigra pars reticulata (SNr), and it sends dense topographical projections throughout the striatum. It is also reciprocally connected with the cortex, as part of a collection of parallel cortical-basal-thalamic loops (Berendse & Groenewegen, 1990; Deniau et al., 1992; Foster et al., 2021; Gonzalo-Martín et al., 2024; Grillner, 2025; Lee et al., 2020; Mandelbaum et al., 2019; Redgrave et al., 2010; Schulz et al., 2009; Smith et al., 2014; Stayte et al., 2021; Yan et al., 2008). Three different components of the PF calcium activity we report imply interesting possible circuit-level control that will be important to explore in future studies. First, we saw a sharp increase in PF activity at the onset and offset of the lever cue in our conditioning task, which corresponded with onset of cue-directed orientations. These bursts in PF activity may be supported by excitatory inputs from the superior colliculus (Krout et al., 2001; McHaffie et al., 2005; Redgrave et al., 2010; Schulz et al., 2009) and brainstem cholinergic nuclei (Barroso-Chinea et al., 2011; Huerta-Ocampo et al., 2020; Varela, 2014; Yan et al., 2008), which have established roles in directing head and eye movements in response to activating sensory information (Arber & Costa, 2022; Hikosaka et al., 2006; Poisson et al., 2025).

Second, sustained PF neuron activity developed across cue learning, with elevation above baseline for the entire duration of the cue period. We speculate that this sustained signal could be related to plasticity of cortical loop connections (Gonzalo-Martín et al., 2024; Mandelbaum et al., 2019), or disinhibition of SNr due to enhanced dorsolateral striatum activity as habitual behavior emerges (Bornhoft et al., 2025; Grillner, 2025; Lee et al., 2020; Mohebi et al., 2024; Yin et al., 2009). In this manner, the PF may integrate nonspecific “bottom up” brainstem attentional signals about salient events with “top down” cortical-basal-ganglia loop signals based on learning, to allow for appropriate actions (Kimura et al., 2004). During extinction, sustained cue-related PF activity diminished, but large signal increases persisted at cue onset and offset. These peaks could be attributed to redirecting attention in response to the changing sensory state of the environment, in service of exploration and behavioral flexibility. This interpretation is consistent with the PF’s role in reversal learning and changing contingencies (Bradfield & Balleine, 2017; Brown et al., 2010; Delacour, 1969; Kato et al., 2011; Minamimoto & Kimura, 2002), which is also heavily dependent on cortex (Izquierdo et al., 2017; Klein-Flügge et al., 2022; Stayte et al., 2021).

Third, we show that reward consumption behavior was associated with a pause in PF activity. This, along with the lack of intracranial self-stimulation behavior we observed for PF neuron activation, suggests that the PF does not signal the inherent value of a stimulus. Given that PF activity encoded cue-related orientations and drove turning behavior, we predict that the pause during reward receipt is important for maintaining stable head position during the time when it would be disadvantageous to shift attention. We also found that repeated activation of neurons in the PF potentiated head/body turning over time, suggesting that engagement of this system supports motor learning to strengthen certain head repositioning motor plans. Collectively, our data motivate a model where dynamic burst-pause activity patterns in the PF could promote gaze adjustments for sensation, exploration, and learning, or stable head positioning, for interface with proximal rewards. The mechanism behind PF inhibition during reward consumption is unclear, but could relate to input from SNr GABAergic neurons. SNr neurons have activity time-locked to actions (Fan et al., 2012), are modulated by reward-predictive cues (Sato & Hikosaka, 2002) and reflect head position (Barter et al., 2015). They may act as “brake” on head reorientation during reward consumption, in a similar fashion to saccade, licking, and reaching control (Arber & Costa, 2022; Chevalier & Deniau, 1984; Falasconi et al., 2025; Hikosaka et al., 2006; Rossi et al., 2016). SNr neurons also project to brainstem areas that input to the PF, which could provide a disynaptic braking mechanism (Arber & Costa, 2022; McElvain et al., 2021).

Our results, based on population level recordings of the PF, suggest that dynamic encoding of attentional shifts that scale with learning is a relatively prominent feature among PF neurons. Notably, previous studies indicate that not all PF neurons are equally engaged by head movements, with subpopulations encoding different directions of head velocity, and others with relatively movement insensitive activity profiles (Fallon et al., 2023; Watson et al., 2021; Y. Zhang et al., 2022). Critically, even in our population level signals, we observed PF activity during cue presentations when rats were moving and also in response to the cue when rats were stationary, suggesting that the PF multiplexes information related to both movement and attention or sensation. Further studies are needed to determine if specific populations of PF neurons are engaged to signal different features of cue-guided learning in our task, and the extent to which heterogeneous activity profiles map onto PF output projections and/or other anatomical features.

Given that PF neurons project to a diverse array of cortical, striatal, and subthalamic nuclei, we would expect that distinct features of reward seeking, such as cue detection, orientation/movement, and action-outcome associations may be encoded by specific PF input-output channels (Gonzalo-Martín et al., 2024; Mandelbaum et al., 2019; Watson et al., 2021; Y. Zhang et al., 2022). Our population recordings provide context for future experiments targeting specific PF populations based on connectivity. In particular, striatal PF outputs, which target cholinergic interneurons (CINs) (Huang et al., 2023; Matsumoto et al., 2001; Parent & Descarries, 2008; Yamanaka et al., 2018) may be critical for cue-reward learning and targeted approach, given that striatal CINs are necessary for associating environmental cues with rewards (Collins et al., 2019; Skirzewski et al., 2022). PF inputs to CINs can drive their characteristic multiphasic firing thought to be permissive of dopamine-mediated striatal plasticity (Becchi et al., 2023; Brown et al., 2010; Chantranupong et al., 2023; Ding et al., 2010; Jang et al., 2026; Krok et al., 2023; Mamaligas et al., 2019; Matsumoto et al., 2001; Miller-Hansen et al., 2026; Reynolds et al., 2022; Y.-F. Zhang et al., 2018). Over-exciting CINs via PF input alters their normal firing (Doig et al., 2014), and impairs extinction learning (Huang et al., 2023). While it remains unclear how PF projections to striatum are engaged in a learning context to affect cue orientation and approach behavior, our results support the notion that PF neuron activity patterns are well-timed to be a key player in regulating striatal heterogeneity to support motivation, learning and cue-guided behavior (Bender et al., 2024; Bornhoft et al., 2025; Collins & Saunders, 2020; Cox & Witten, 2019; Parker et al., 2016; Willuhn et al., 2012; Yin et al., 2009).

Our studies identify the PF thalamus as an neural interface between sensory attention and directed action that is important for associative learning. Given the position of this region within cortical and basal ganglia systems fundamental to behavioral control, dysregulation of normal activity in the PF likely contributes to a broad array of motivational, cognitive, and motor diseases.

## METHODS

### Subjects

Male and female Long-Evans rats (N=32, 17M, 19F) were used, 3-6 months old with starting weights of 200-500 g. Rats were housed in ventilated cages in a vivarium with a 12 hour light:dark cycle. All procedures involving animal subjects were in compliance with Institutional Animal Care and Use Committee (IACUC) approval and in accordance with the National Institutes of Health’s animal care guidelines.

### Stereotaxic surgery

Rats were induced under 5% isoflurane anesthesia, and placed in a stereotaxic frame fitted with a nose cone to maintain at 1–3% anesthesia throughout surgery. At the beginning of surgery, rats received carprofen (5 mg/kg), cefazolin (70 mg/kg), and saline subcutaneously. Rats received an incision to reveal their skull, which was leveled and holes were made for viral injections (-4.2 mm Posterior, +/-1.2 ML, -6.2 Ventral, 250 nL), and implants relative to bregma and skull surface, along with 4 holes for screws to secure headcaps. To target PF neurons for optogenetic activation, we expressed AAV5-hsyn-hChR2-EYFP (250 nL, Addgene #26973; initial titer 7×10^12 vg/ml). For PF neuron calcium recordings, we expressed AAV5-hsyn-GCaMP8f (250 nL, Addgene #162376 1×10^13 vg/ml). Viral injections were made via Hamilton syringe at 100 nL/min. Post injection, the syringe was left in the brain 200 um above the injection site for 10 minutes to allow the virus to settle. Four screws were attached to the skull, and a headcap made of dental cement was affixed around an implant for photometry (-4.2 Posterior, +/-1.2 ML, -6.0 Ventral, 9 mm length, 400 µm diameter, Doric Lenses) or optogenetics (-4.2 Posterior, +/-1.2 ML, -5.7 Ventral, 10mm length, 300μm diameter, Thorlabs). At the end of surgery, rats were treated with lidocaine and anti-bacterial ointments and monitored on a heating pad post-anesthesia until they were alert and awake. Rats were given carprofen and cefazolin for 3 days post surgery and monitored for 7 days post surgery for weight and visual signs of distress. Photometry recordings and optogenetic manipulations commenced at least 3 weeks after surgery to allow for adequate recovery and viral expression.

### Behavioral training

Behavioral training procedures were based on established pavlovian conditioned approach methods (Flagel et al., 2011; Saunders & Robinson, 2012). Rats were mildly food deprived starting one week before behavioral procedures, and maintained at 90% free feeding weight for the duration of the experiment. All behavior took place in soundproofed operant chambers, outfitted with two retractable levers, lights, fan, and two nose pokes with three cue interior cue lights (Med Associates). A port for reward delivery was positioned between the levers, equipped with an infrared beam to detect port entry and exit. Chambers had a hole in the top for a flexible fiber patch cord for fiber photometry recording, as well as a flexible arm with commutator for an armored cable for laser stimulation. For all stages of behavior, lights and a fan were left on throughout the session. Behavior boxes were wiped with ethanol and bedding was replaced between each rat.

### Magazine training

Prior to conditioning, all rats were habituated to cable tethering and Med Associates chambers while they received 30 non-contingent 15% sucrose rewards at random intervals with chamber lights and fans running. Rats that had low consumption of sucrose in the chamber were given 1-2 days of standard chow soaked in sucrose.

### Cue habituation

Rats (n=7) received 2 cue habituation sessions wherein 10 8 sec lever presentations were delivered on a variable interval averaging 77.5 secs (range 50”-105”).

### Reward conditioning

Cohorts for both photometry and optogenetics were reward conditioned for 12-15 days. Each session consisted of 40 cue presentations with a variable intertrial interval averaging 77.5 secs (range 50-105 s). In each trial, the cue (lever) was extended for 8 seconds. Concurrent with retraction of the lever, a 3 second 15% sucrose reward was delivered into the magazine via a pump. Acquisition of learning was measured via duration of time in the reward port during the cue.

### Extinction

Rats (n=8) went through extinction for 2 to 3 days. Each session of extinction was identical to reward conditioning, except for no reward was delivered at cue offset.

### Intracranial self-stimulation

After conditioning, optogenetics rats (n=11) underwent three one-hour sessions of intracranial self-stimulation (ICSS). During ICSS, rats had the choice between an active nose poke, designated by three interior cue lights that flashed with each poke, and an inactive nose poke without cue lights. Each active poke triggered a 1-second 20Hz laser stimulation, and each inactive poke had no consequence.

### Optogenetics

For optogenetic experiments, rats were tethered on all experimental days to a fiber optic cable, armored for flexibility and light protection on a flexible arm fitted with a commutator. On stimulation days, they received 5ms pulses of a 473 nm laser (OptoEngine) at 20 Hz frequency for 1-5 seconds. Laser power was set to 10 mW prior to each stimulation session, measured at constant power, resulting in individual pulses of ∼2 mW. Stimulations were time-locked to MedPC TTL events.

#### Reward conditioning

Rats received 5-second stimulations time-locked to cue onset (paired), or within the inter-trial interval (unpaired), on 50% of trials. Stimulations occurred on days 1, 5, 7, and 12 of training.

#### Intracranial self-stimulation

Rats received 1 sec stimulations time-locked with onset of active nose pokes.

### Fiber photometry recordings

To assess calcium dynamics in PF neurons during pavlovian conditioning, we measured GCaMP fluorescence using fiber photometry. For all behavioral experiments, rats were tethered to a low autofluorescence optic cable sheathed in a lightweight armored jacket (Doric-Lenses). Recordings were performed every other day to avoid photobleaching GCaMP fluorescent proteins. Recording data included here are sampled from the early (Day 1), middle (Day 3), and late (Day 6, 7, or 8) of reward conditioning. On recording days, 415 nm (isosbestic) and 465 nm (Ca2+ dependent) LEDs (Doric-Lenses) set at 50 µW were delivered via a fluorescence mini-cube (Doric-Lenses), and modulated at 211 Hz and 330 Hz, respectively. Fluorescence from the implanted optic fiber was transmitted through the mini-cube to be filtered, amplified, and focused onto a high-sensitivity photoreceiver (Newport, Model 2151). A real-time signal processor (RZ5P, Tucker Davis Technologies) using Synapse software modulated the power of the LED output and recorded isosbestic (Ca2+ independent) and Ca2+ dependent signals, sampled at 6.1 kHz. Events during the behavioral session (e.g. cue and reward presentations) were recorded in the photometry data file with a TTL signal time stamp from the MED-PC behavioral program. Videos of each behavioral session were recorded at 10–20 FPS with corresponding timestamps of each frame in the photometry file.

### Fiber photometry analysis

Recordings were analyzed with a MATLAB (Mathworks) pipeline. First, signals were low-pass filtered at 6 Hz, downsampled to 40 Hz, and a least squares linear fit was applied to the isosbestic channel (415 nm) to align it to the calcium-dependent, 465 nm, channel. The fitted isosbestic channel was then used to normalize the 465 signal with the calculation ΔF/F = (465-nm signal—fitted 415-nm signal)/(fitted 415 nm signal). Individual trial traces were z-scored to the 2 s immediately preceding cue and port entry events in order to avoid effects of drift across the session. Area under the curve (AUC) values took the z-scored trace and calculated numerical integration via the trapezoidal method using the trapz function (MATLAB). AUC and peak Z-score values were calculated for event windows of interest.

### Automated pose estimation

Markerless tracking of rat body parts was conducted using version 2.2.1.1 of the DeepLabCut (DLC) Toolbox (Mathis et al., 2018), and analysis of movement features based on these tracked coordinates was conducted in MATLAB. DeepLabCut 2.2.1.1 was installed in an Anaconda environment with Python 3.8.4, CUDA 11.7 and Tensorflow 2.10. DeepLabCut Model: 2090 frames from 35 videos (32 different rats, 3 experiments) were labeled, and 807 outlier frames were re-labeled to refine the network. Labeled frames were split into a training set (95% of frames) and a test set (5% of frames). A ResNet-50-based neural network (Insafutdinov et al., 2016) was used for 1,030,000 training iterations. After the final refinement, we found the test error was 4.1 pixels, the training error was 3.13 pixels and with a p-cutoff of 0.85 training error was 2.99 pixels, and the test error was 3.68 pixels. The body parts labeled included the nose, eyes, ears, fiber optic implant, shoulders, tail base, and an additional three points along the spine. Features of the environment were also labeled, including the 4 corners of the apparatus floor, two nose ports, two cue lights, two reward ports, and 3 LED indicator lights when active. For optogenetics experiments, LED cue lights were time-locked to events such as cue onset and laser stimulations, and were used to sync video recordings with events. DLC coordinates and confidence values for each body part and frame were imported to Matlab and filtered to exclude body parts/features from any frame where the confidence was <0.7. For labeled features of the environment, which have a fixed location, the average coordinates for that recording were used for analysis. To convert pixel distances to the real chamber dimensions, for each video, a pixel-to-cm conversion rate was determined. The distance (in pixels) between each edge of the environment floor and the diagonal measurements from corner to corner were measured, and these values were divided by the actual distance in cm. The mean of these values was then used as the conversion factor.

#### Movement speed

was calculated from the implant coordinates frame by frame using the formula: [distance moved (pixels per cm) * framerate] to give movement speed in cm/s.

#### Cumulative angle

was calculated by adding the change in the angle (degrees) between the vector between the shoulder point and the mid back point on the current frame and this same vector in the previous frame using the formula angle = atan2(norm[cross(a,b)], dot(a,b)). Resulting angle values were normalized for the implant hemisphere so that increases in the cumulative angle reflected the movement direction (ipsilateral to the implant).

#### Locomotion bouts

were identified using the speed of the implant, midback, and shoulder body parts. Bouts were counted when all visible body parts were above their corresponding movement threshold for any single video frame, while the transitions in and out of a movement bout were demarcated by a sliding window when the speed of 2 or more body parts exceeded the detection threshold for 0.3 s. Movement thresholds used a scale factor based on rat size and were set separately for body parts on the head to account for the finer movements relative to the body. Movement periods needed to be 0.5s or longer to be considered as a bout. Angular velocity was calculated as the absolute change in degrees/sec of a deflection of a vector from the rats’ implant to mid back.

#### Orientations

were defined as epochs when a vector from the rat’s implant to mid back was within 30 degrees or less of the reward magazine for a minimum of 0.4 seconds. A sliding window of 0.2 s was used to identify transitions in and out of orientation epochs.

### Histology

At the end of experiments, rats were injected with Fatal Plus pentobarbital (2 ml/kg; Paterson Veterinary) before being transcranially perfused with cold 1X PBS and 4% paraformaldehyde. Brains were extracted and left in 4% paraformaldehyde for 24 hours before being preserved in 50% sucrose in PBS for at least 48 hours. Brains were sliced at 50uM in free floating sections and mounted onto glass slides. Tissue was dried before being coverslipped with Vectashield antifade with DAPI mounting media and sealed with clear nail polish as needed. Viral targeting and damage were assessed through imaging on a flourescent microscope (Keyence BZ-X710) with a 4X and 10X air immersion objective. Images were compared to a rat brain atlas (George Paxinos, 2007).

### Statistical analysis

All statistical analyses were performed within Graphpad prism software. Analyses used were 2-way ANOVA (including mixed models) and 1-way ANOVA. Bonferroni-corrected post hoc comparisons were used to compare groups. Orientation bias distributions were compared using the Kolmogorov-Smirnov test. All statistical tests were two-tailed. Data are expressed as subject mean +/-SEM. Statistical significance was set to p <0.05.

## Funding and Acknowledgements

The authors have no biomedical financial interest or other conflicts of interest. This work was supported by a MnDrive Neuromodulation fellowship from the University of Minnesota (LCK) and NIH grants R01 MH129370, R01 MH129320, and R01 DA057292 (BTS). The authors thank all members of the Saunders labs for support and feedback.

## Supplemental Material

**Fig S1.**
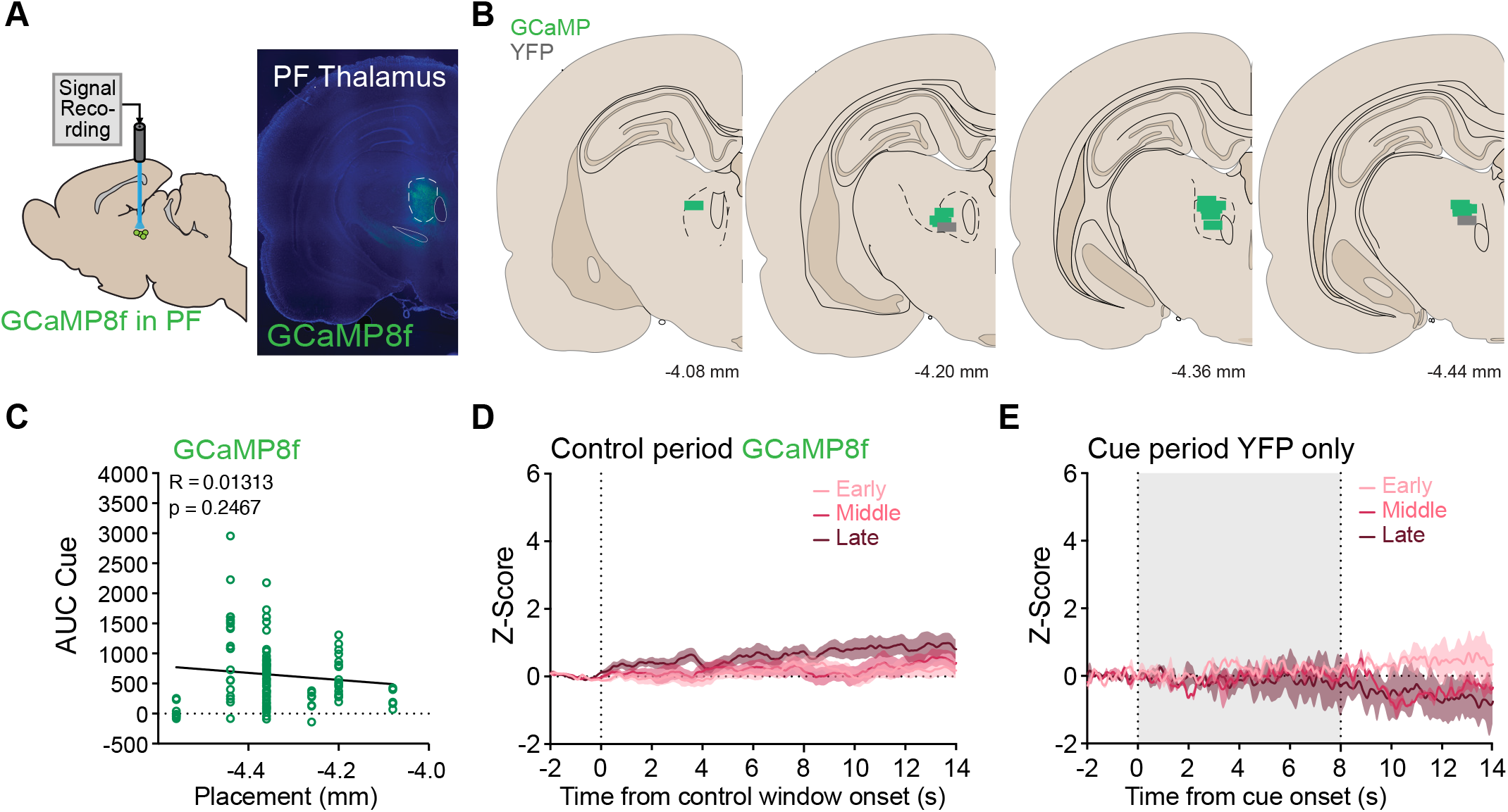
Fiber photometry recording histology and control signals. A) Representative image of GCaMP expression and fiber placement. B) Fiber placements in the parafasicicular (PF) thalamus. C) We found no correlation between recording location along the anterior-posterior axis and the magnitude of GCaMP cue responses (R=0.01313, p=0.2467). D) Recordings of PF GCaMP during control windows (within the inter-trial interval) showed no changes. E) Control photometry recordings made from a YFP-only virus showed no change in signal during cue presentations. Data shown reflect mean +/-SEM.

**Fig S2.**
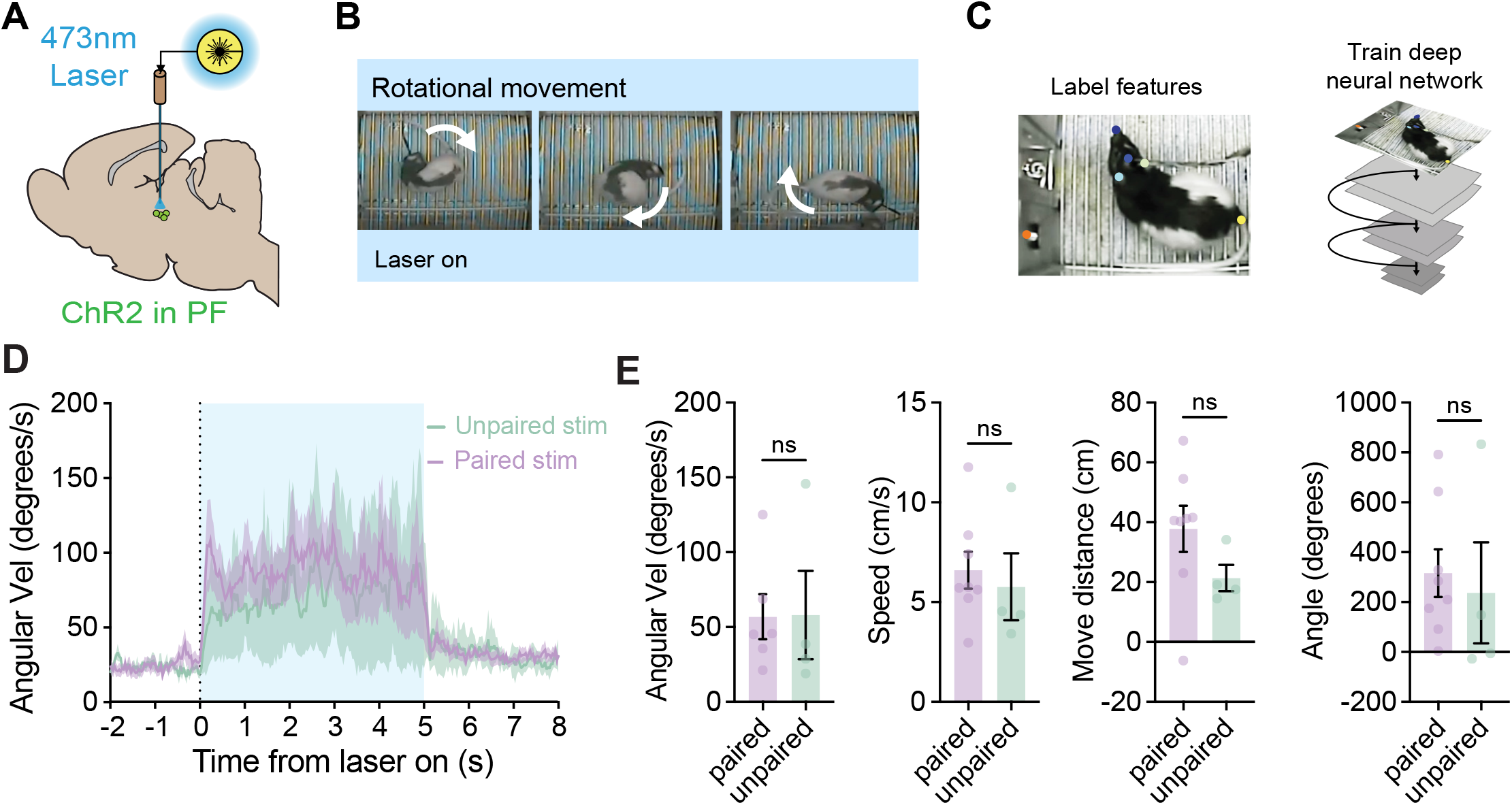
Turning behavior during unpaired PF activation. A) ChR2-YFP was expressed and optic fibers were implanted unilaterally in the PF thalamus. Optogenetic stimulation (20 Hz, 5 sec) was delivered either during the cue period (paired, n=8) or inter-trial-interval (unpaired, n=4). B) Representative images of rotational movement while the laser was on. Rats curled and turned their entire body in circles, directed by their head. C) Schematic of behavioral analysis via DeepLabCut. D) Average angular velocity on the last day for paired and unpaired stimulation conditions. E) There were no significant differences in average angular velocity, average speed, move distance, or cumulative angle between paired stimulation and unpaired stimulation groups. Data shown reflect mean +/-SEM.

**Fig S3.**
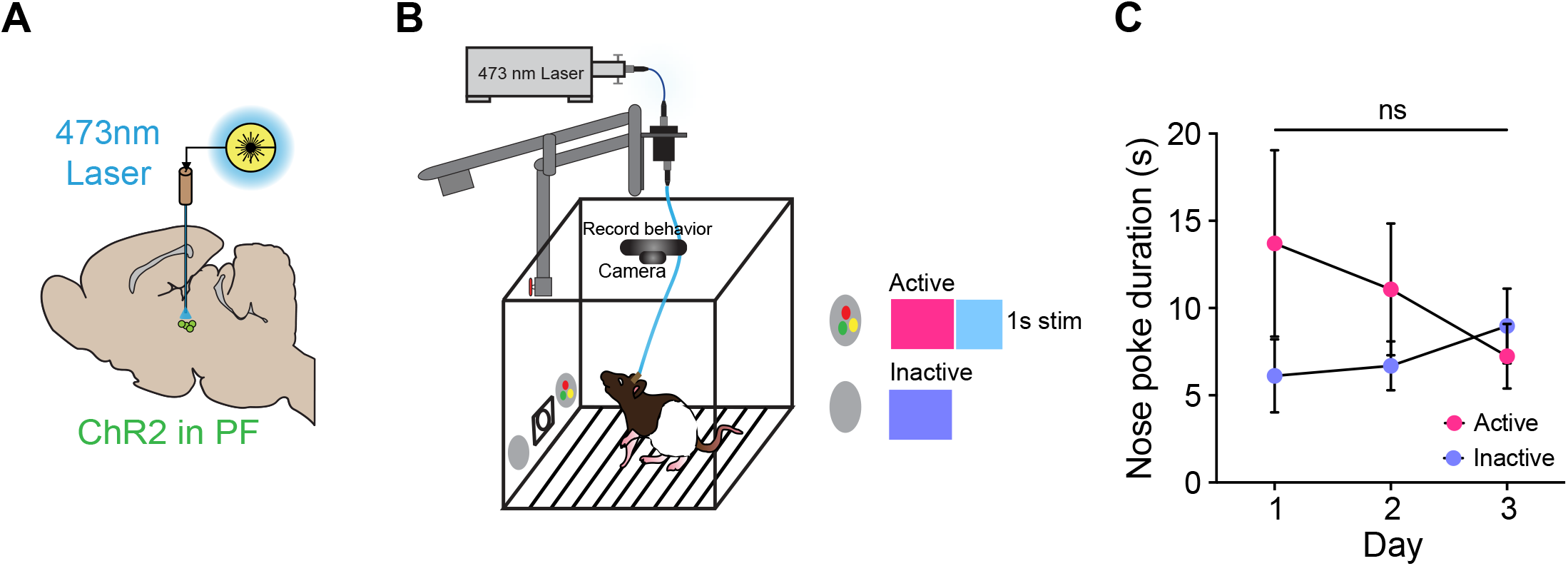
PF stimulation is not reinforcing. A) ChR2 was expressed unilaterally in the PF thalamus. B) Rats (n=11) were given the opportunity to self-stimulate the PF, where active nose pokes coincided with a brief visual cue inside the poke and a 1-s 20 Hz laser pulse. C) There was no difference in nose poke duration between active and inactive nose pokes. Data shown reflect mean +/-SEM.

